# Single synapse indicators of impaired glutamate clearance derived from fast iGlu*_u_* imaging of cortical afferents in the striatum of normal and Huntington (Q175) mice

**DOI:** 10.1101/455758

**Authors:** Anton Dvorzhak, Nordine Helassa, Katalin Török, Dietmar Schmitz, Rosemarie Grantyn

## Abstract

Changes in the balance between glutamate (Glu) release and uptake may stimulate synaptic reorganization and even synapse loss. In the case of neurodegeneration, a mismatch between astroglial Glu uptake and presynaptic Glu release could be detected if both parameters were assessed independently and at a single synapse level. This has now become possible due to a new imaging assay with the genetically encoded ultrafast Glu sensor iGlu*_u_*. We report findings from individual corticostriatal synapses in acute slices prepared from mice aged >1 year. Contrasting patterns of short-term plasticity and a size criterion identified 2 classes of terminals, presumably corresponding to the previously defined IT and PT synapses. The latter exhibited a higher degree of frequency potentiation/residual Glu accumulation and were selected for our first iGlu*_u_* single synapse study in Q175 mice, a model of Huntington’s disease (HD). It was found that in HD the time constant of perisynaptic [Glu] decay (TauD, as indicator of uptake) and the peak iGlu*_u_* amplitude (as indicator of release) were prolonged and reduced, respectively. Treatment of WT preparations with the astrocytic Glu uptake blocker TFB-TBOA (100 nM) mimicked the TauD changes in homozygotes (HOM). Considering the largest TauD values encountered in WT, about 40% of PT terminals tested in Q175 heterozygotes (HET) can be classified as dysfunctional. Moreover, HD but not WT synapses exhibited a positive correlation between TauD and the peak amplitude of iGlu*_u_*. Finally, EAAT2 immunoreactivity was reduced next to corticostriatal terminals. Thus, astrocytic Glu transport remains a promising target for therapeutic intervention.

**SIGNIFICANCE STATEMENT:** Alterations in astrocytic Glu uptake can play a role in synaptic plasticity and neurodegeneration. Until now, sensitivity of synaptic responses to pharmacological transport block and the resulting activation of NMDA receptors were regarded as reliable evidence for a mismatch between synaptic uptake and release. But the latter parameters are interdependent. Using a new genetically encoded sensor to monitor [Glu] at individual corticostriatal synapses we can now quantify the time constant of perisynaptic [Glu] decay (as indicator of uptake) and the maximal [Glu] elevation next to the active zone (as indicator of Glu release). The results provide a positive answer to the hitherto unresolved question whether neurodegeneration (e.g. Huntington’s disease) associates with a glutamate uptake deficit at tripartite excitatory synapses.

## INTRODUCTION

A low level of steady state glutamate concentration [Glu] is an important prerequisite for high spatial and temporal discrimination of afferent signals. Electrogenic transport of Glu from the environment of active synaptic terminals into astroglial cells secures resting [Glu] levels below 100 nm (Bergles et al., 1999; Marcaggi and Attwell, 2004; Tzingounis and Wadiche, 2007; Nedergaard and Verkhratsky, 2012; Papouin et al., 2017; Rose et al., 2018). Compared to other synaptically enriched proteins, including the AMPA receptors, excitatory amino acid transport proteins (EAAT1 and EAAT2, GLAST and GLT1 in rodents) are very abundant (Lehre and Danbolt, 1998; Marcaggi and Attwell, 2004; Cahoy et al., 2008) forming clusters on the perisynaptic astroglial processes (PAPs) next to the sites of transmitter release (Lehre and Danbolt, 1998; Melone et al., 2011). However, the proximity between the sites of synaptic Glu release and astrocytic uptake could vary according to the type of synapse or the functional state of the involved cells (Octeau et al., 2018).

Insufficient expression/activity of EAAT2 is considered among the mechanisms promoting excitotoxic damage and neurodegeneration (Pekny et al., 2016; Verkhratsky et al., 2016). As for HD, there is full agreement that transcription of the EAAT2 encoding gene *SLC1A2* and tissue uptake of radio-labelled EAAT2 substrates are reduced in comparison with healthy controls (Lievens et al., 2001; Behrens et al., 2002; Shin et al., 2005; Miller et al., 2008; Bradford et al., 2009; Faideau et al., 2010; Huang et al., 2010; Menalled et al., 2012; Grewer et al., 2014; Meunier et al., 2016). Records from striatal astrocytes (Dvorzhak et al., 2016) suggested a 20-30% decrease of the glutamate uptake activity in two mouse models of HD, R6/2 and Q175. Yet some caution is needed as it has remained unclear whether or not the well-documented reduction of astrocytic Glu transport is to be regarded as a primary cause of synapse dysfunction/loss, or merely as an epiphenomenon reflecting glial adjustment to the massive pruning of glutamatergic terminals for other yet unknown reasons (Deng et al., 2013; Rothe et al., 2015). As synapse degeneration is likely to progress in a rather asynchronous manner resulting in a co-existence of dysfunctional and more or less healthy terminals, a satisfying answer regarding the adequate performance of astrocytic glutamate uptake in HD can only be obtained at the single synapse level and under consideration of the individual uptake/release relationships (Barbour, 2001; Nahir and Jahr, 2013; Jensen et al., 2017; Reynolds et al., 2018).

The striatum as the most affected brain structure in HD (Khakh et al., 2017) is well suited for selective activation of glutamatergic synapses as it lacks intrinsic glutamatergic neurons. Glutamatergic afferents originate in the medial thalamus and the cerebral cortex (see (Reiner and Deng, 2018) for more). Corticostriatal connections are formed by at least two distinct populations of pyramidal neurons, localized in layers 2/3 and 5. The axons originating in layer 2/3 establish bilateral intra-telencephalic (IT) connections, while layer 5 axons enter the pyramidal tract (PT) and lack telencephalic collaterals to the contralateral side. Elegant electrophysiology (Kincaid et al., 1998) and electron microscopy studies (Reiner et al., 2010) discovered a number of differences between PT and IT afferents and their synaptic varicosities. In view of this diversity one could expect some type-dependent differences in the release characteristics and, accordingly, differential sensitivity to factors that may cause an uptake/release imbalance.

Here we report the results of the first single synapse experiments in striatal slices of adult mice performed with the ultrafast Glu sensor iGlu*_u_* (Helassa et al., 2018). We have addressed three main questions: 1) How does activation frequency affect the Glu release and uptake at corticostriatal terminals? 2) Does HD produce an uptake/release mismatch? 3) If so, could the indicators of uptake and/or release be used to identify dysfunctional synapses?

## MATERIALS AND METHODS

### Animals

The work described here has been carried out in accordance with the EU Directive *2010/63/EU* for animal experiments and was registered at the Berlin Office of Health Protection and Technical Safety (G0233/14 and G0218/17). Z-Q175-KI mice were obtained from CHDI (“Cure Huntington’s Disease Initiative”, see stock # 027410 of the Jackson Laboratory, Bar Harbor, USA). The number of CAG repeats ranged from 182 to 193. The recordings were performed in animals of either sex at an age of 51 to 76 weeks.

### Plasmids

pAAV-CaMKIIa-ChR2(H134R)-EYFP and pAAV-CaMKIIa-hChR2(E123 T/T159C)-EYFP (Addgene, Watertown, USA #26969 and #35511) were gifts from Karl Deisseroth. To create pAAV-CaMKIIa-iGlu*_u_* (Addgene #75443), the iGlu*_u_* gene was amplified by PCR from pCI-syn-iGlu*_u_* (Addgene #106122) using Phusion polymerase (forward 5’-CATCAGGATCCATGGAGACAGACACACTCC-3’, reverse 5’-GTATGGAATTCCTAACGTGGCTTCTTCTGCC-3’) and cloned into pAAV-CaMKIIa-hChR2(H134R)-EYFP by restriction-ligation using BamHI/EcoRI restriction enzymes (NEB) and T4 DNA ligase (NEB). AAV9-CaMKIIa.iGlu*_u_*.WPRE-hGH and AAV9-CaMKIIa.hChR2(E123 T/T159C)-EYFP.hGH were packaged at University of Pennsylvania Vector Core (Penn Vector Core).

### Drugs and antibodies

All substances were obtained from Sigma Aldrich/Merck, Taufkirchen, Germany, except TTX (Abcam, Cambridge, UK) and TFB-TBOA (Tocris, Bristol, UK). The primary antibodies included those to vGluT1 (1:1000, guinea pig, Synaptic Systems #135304) and EAAT2 (1:2000, rabbit, Abcam #ab41621). Secondary antibodies against guinea pig and rabbit, were conjugated to Alexa 488 or 555 and obtained from Life Technologies (#A-11073, #A-21429, respectively).

### Injections and brain slice preparation

The animals were anesthetized by intraperitoneal injection of a mixture containing 87.5 mg/kg ketamine and 12.5 mg/kg xylasine before receiving 4 intracortical injections of AAV9-CamKII.iGlu*_u_*.WPRE-hGH (7.34*10^13^gcC/ml −0.3 µl) or 1 intracortical injection of AAV9-CaMKIIa.hChR2(E123 T/T159C)-EYFP.hGH (6.28*10^12^ gc/ml −1 µl) at the following coordinates with respect to bregma (mm): anterior 1.5, lateral 1.56, 1.8, 2.04, 2.28 and ventral 1.7. About 3 weeks (ChR2) or 6 weeks (iGlu*_u_*) later the animals were anesthetized with isoflurane, transcardially perfused with cooled aerated saline containing (in mM): N-methylglucamine chloride (NMDG) 92, KCl 2.5, NaH_2_PO_4_ 1.25, NaHCO_3_ 25, glucose 20, CaCl_2_ 0.5, MgCl_2_ 10, sodium pyruvate 3, and sodium ascorbate 5 (pH 7.35, 303 mosmol/l). After decapitation and removal of the brains, parasagittal (10 deg off) sections (300 µm) containing the striatum were prepared as previously described (Dvorzhak et al., 2016). The slices were kept in artificial cerebrospinal fluid (ACSF) containing (in mM): NaCl 125, KCl 3, NaH_2_PO_4_ 1.25, NaHCO_3_ 25, CaCl_2_ 2, MgCl_2_ 1, glucose 10 (pH 7.3, 303 mosmol/l), supplemented with (in mM): sodium pyruvate 0.5, sodium ascorbate 2.8 and glutathione 0.005. These perfusion and recovery solutions preserved the astrocytes better than physiological ACSF, sucrose- or choline-containing solutions, the criteria being resting membrane potential at break-in (WT: <=-85 mV). Q175 HOM were also injected with CEF (5 consecutive days before testing, 200 mg/kg i.p.) or the respective control solution (physiological saline).

### Quantification of synaptic [*Glu*] elevations with iGlu_*u*_

The biophysical characteristics of the new ultrafast Glu sensor (iGlu*_u_*) were already described (Helassa et al., 2018). Briefly, responses to saturating Glu concentration (10 mM) were recorded in transduced HEK293 T cells. An iGlu*_u_* off rate of 2.1 ms was determined using recombinant purified protein and stopped flow fluorimetry. For the imaging of synaptically released Glu, slices were submerged into a perfusion chamber with a constant flow of oxygenated ACSF at a rate of 1-2 ml/min. Temperature during the recordings was maintained at 26 - 27 °C. In non-stimulated acute slices from >1 year old mice corticostriatal varicosities were visualized in the dorsal striatum using a Zeiss W Plan-Apochromat 63x /NA 1.0 water immersion objective and brief (180 ms) discontinuous exposure to a 473 nm laser beam focused to a circular area of ∼4.5 µm in diameter centered to a presynaptic varicosity. The distance to the nearest other fluorescent varicosity was typically 3 to 5 µm. The size of non-stimulated boutons was derived from the area of supra-threshold pixels, the threshold being defined as mean ROI intensity + 3 SD. For evaluation of evoked responses, the iGlu*_u_* fluorescence was acquired at a frequency of 2.5 kHz from a rectangular ROI of 4 µm x 4 µm (20 × 20 pixels, binning 2) using a sCMOS camera (Andor Zyla4.2 plus) attached to a Zeiss wide field microscope (AxioObserver). In-house written software routines controlled the laser, camera and electrical stimulation of the axon/bouton. Each pixel of the ROI was evaluated separately for the construction of time- and space-dependent [Glu] profiles after release. The iGlu*_u_* intensity values were expressed as supra-threshold pixel fluorescence ΔF in % of the mean baseline fluorescence derived from the data points acquired during a 50 ms period prior to stimulation. The stimulus-induced changes of suprathreshold ΔF/F in time or space are referred to as “iGlu*_u_* transients” or simply “transients”.

For the quantification of iGlu*_u_* at single synapses we defined the following key parameters. The boundaries of the presynaptic bouton at rest (prior to any stimulation) were calculated from the F values at rest and included pixels with F larger ROI mean + 3 SD (see Fig. 1D, area outlined in blue). The area of suprathreshold pixels at rest was approximated as a circle, and the resulting virtual “*Bouton diameter*” was used as indicator of bouton size. The term “*Peak amplitude*” refers to the peak ΔF/F value of an averaged intensity transient derived from all suprathreshold pixels (see Fig. 1E, distance between dotted red lines). “*Tau decay*” abbreviated as*“TauD”* is the time constant of decay derived by fitting a monoexponential function to the decay from the peak of the averaged transients (see Fig. 1E, amplitude between dotted red lines). The spatial extension of the iGlu*_u_* signal is described on the basis of a virtual diameter derived from the area of all suprathreshold pixels combined to form a virtual circle. The respective diameter is referred to as “*Spread*”. The term “*Peak* s*pread*” refers to the peak value of the averaged spread transient (see Fig. 1F, difference between dotted red lines). The indicator “*Residual ΔF/F*” is derived from fitting a double exponential function to the iGlu*_u_*transient after the last stimulus. It corresponds to the ΔF/F value at the intercept between the fast and slow phase of iGlu*_u_*decay (see Fig. 2E, red horizontal line). “*Integral ΔF/F*” refers to the sum of all responses during a series of 6 stimuli at 100 Hz within a period of 70 ms starting with the first stimulus. Dysfunctional synapses could best be detected by analysis of single pixel iGlu*_u_* using the pixel with the highest iGlu*_u_*elevation at any given terminal. The highest iGlu*_u_* elevations were always found within or next to the bouton at rest. The peak amplitude of the single pixel transient with the highest iGlu*_u_* elevation will be referred to as “*Maximal amplitude*” (see Fig. 4A-C, difference between red dotted lines). The respective TauD values are referred to as “*TauDmax*”. In the following text, these parameter names will be written in italics and capitals to underline that these are pre-defined indicators introduced for the convenience of the present single synapse analysis.

**Fig. 1.**
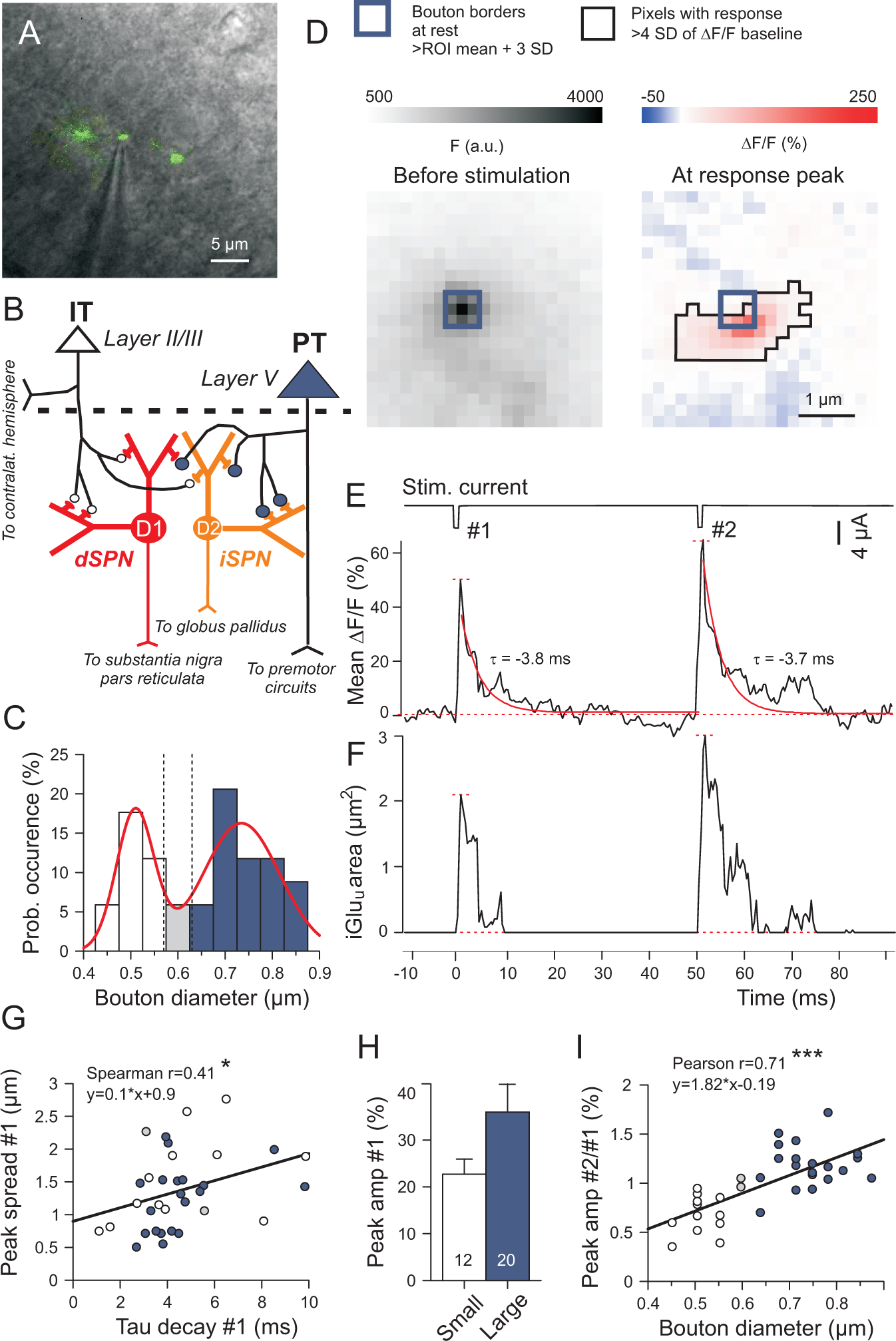
Monitoring single synapse Glu transients in acute slices from adult mice after expression of the genetically encoded ultrafast Glu sensor iGlu_u_ in corticostriatal neurons. (A) Resting iGlu_u_ fluorescence merged to the respective 63x DIC image of a corticostriatal slice showing an axon with 3 adjacent varicosities and a stimulation pipette at the central bouton. (B) Simplified scheme of the corticostriatal circuitry (Reiner et al., 2010), illustrating the concept of preferential projection of pyramidal tract (PT) neurons to indirect pathway striatal projection neurons (iSPNs) and intratelencephalic (IT) neurons to direct pathway SPNs (dSPNs), with size-differences between the IT and PT terminals. (C) Bimodal distribution of bouton diameters as determined by the supra-threshold resting fluorescence before stimulation. Boutons with diameter > = 0.63 µm were defined as “Large” and assumed to be issued by PT axons. (D) Example of a type PT bouton with the respective iGlu_u_ fluorescence at rest (left) and at the peak of an AP-mediated iGlu_u_ response (right). (E, F) iGlu_u_ responses recorded from the bouton shown in (A, D). Experiment in 2 mM Ca^2+^ and 1 mM Mg^2+^. (E) Simultaneous recording of stimulation current (upper trace) and mean intensity of supra-threshold pixels (bottom trace). Peak amplitudes (between dotted red horizontal lines) and a monoexponential function fitted to the decay from this peak (red overlay) rendered the TauD values next to the fitting curves. (F) Plot of spread against time (for a definition see Methods). Peak spread: difference between dotted red horizontal lines. (G) Positive correlation between peak spread and TauD after stimulus #1. (H) “Peak amplitude of responses to stimulus #1. There is no difference between small and large terminals. (I) Significant correlation between the PPR of peak amplitudes and bouton diameters. Asterisks denote: * - P < 0.05, ** - P < 0.01, *** - P < 0.001.

**Fig. 2.**
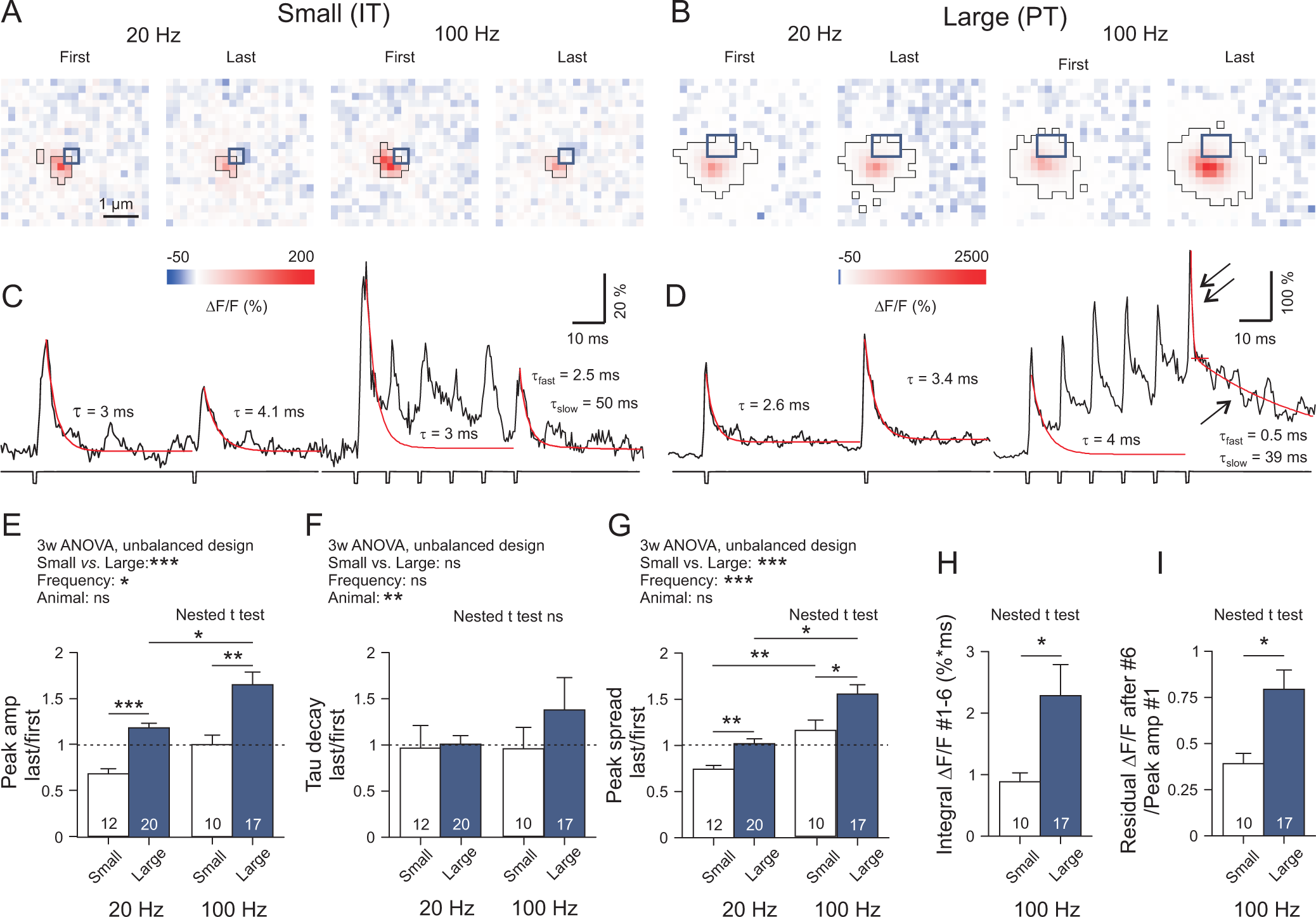
Contrasting dynamics of Glu release from small and large corticostriatal terminals. (A-D) Specimen records of AP-mediated iGlu_u_ signals from small and large boutons, as obtained with 20 and 100 Hz stimulation. All iGlu_u_ transients were elicited in an AP-dependent manner in 2 mM Ca^2+^. Experimental conditions as in Fig. 1. The large terminals produced a significant build-up of residual iGlu_u_ (single arrow), i.e. fluorescence that added to the fast stimulus-locked transients (double arrow). (E-I) Quantification of results. Note that the time integral of all supra-threshold pixel intensities generated by a 6-pulse train at 100 Hz during a sampling period of 70 ms was much bigger in large boutons (I). 3-way ANOVA statistics: E) Leven’s test F(df1 = 29, df2 = 29) = 3.802, P < 0.001. Small vs. Large F(1,29) = 19.507, P < 0.001. Frequency F(1,29) = 7.207, P = 0.012. Animal F(9,29) = 0.765, P = 0.649. Animal-Frequency F(8,29) = 1.154, P = 0.359. Animal-Small/Large F(5,29) = 0.46, P = 0.803. Small/Large-Frequency F(1,29) = 0.000, P = 0.999. F) Leven’s test F(df1 = 29, df2 = 29) = 2.202, P = 0.020. Small vs. Large F(1,29) = 1.207, P = 0.281. Frequency F(1,29) = 1.038, P = 0.317. Animal F(9,29) = 4.330, P = 0.002. Animal-Frequency F(8,29) = 2.398, P = 0.041. Animal-Small/Large F(5,29) = 2.206, P = 0.82. Small/Large-Frequency F(1,29) = 0.305, P = 0.585. G) Leven’s test F(df1 = 29, df2 = 29) = 3.202, P = 0.015. Small vs. Large F(1,29) = 11.226, P = 0.002. Frequency F(1,29) = 21.526, P < 0.001. Animal F(9,29) = 0.840, P = 0.575. Animal-Frequency F(8,29) = 0.895, P = 0.533. Animal-Small/Large F(5,29) = 0.818, P = 0.547. Small/Large-Frequency F(1,29) = 0.155, P = 0.696. H) Nested t test: F(1,12) = 6.05. P = 0.032. (I) Nested t test: F(1,25) = 6.47, P = 0.018. Cohen’s D values. E) Small vs. Large at 20 Hz: D = 2.4, Small vs. Large at 100 Hz: D = 1.3. F) D < 0.4 in all pairs. G) Small vs. Large at 20 Hz: D = 1.4, Small vs. Large at 100 Hz: D = 1.1. (H) D = 0.8. I) D = 1.0. (I) G = 1.0. (H) G = 1.0. The asterisks on the horizontal bars on the graphs denote: significance levels according to the nested t test. * -P < 0.05, ** - P < 0.01, *** - P < 0.001.

### Single axon/bouton activation

To induce the Glu release from individual synaptic boutons under physiological conditions, a depolarizing current pulse was applied through an ACSF-filled glass pipette (tip diameter <1 µM, resistance 10 MOhm) placed next to an axon in close proximity with a fluorescent varicosity. Responses were elicited at minimal intensity at a repetition frequency of 0.1 Hz. They disappeared when the pipette was moved by as little as 5 pixel diameters (1 µm). Single bouton recording of iGlu*_u_* in the presence of TTX was performed in elevated (5 mM) [Ca^2+^]_ec_ using a biphasic stimulation. For more details on single bouton activation and recording of unitary EPSCs see (Kirischuk et al., 1999; Kirischuk et al., 2002) and (Dvorzhak et al., 2013a).

### Patch-clamp recording of unitary EPSCs (uEPSCs)

uEPSCs were recorded in the presence of bicuculline methiodide (BMI), as previously described (Dvorzhak et al., 2013b). Briefly, the intra-pipette solution contained (in mM): cesium methane sulfonate 100, CsCl 50, NaCl 5, CaCl_2_ 0.5, EGTA 2.5, Hepes 25, MgATP 2, GTP 0.3 (pH 7.2). uEPSC were induced via optical activation of APs in hChR2(E123 T/T159C)-EYFP expressing corticostriatal axons. Using the point illumination system UGA-42 of Rapp OptoElectronic, the duration and size of the laser pulse was adjusted to activate a synaptic response with distinct threshold. Stimulation was accepted as minimal if the following criteria were satisfied: (i) uEPSC latency remained stable (fluctuations <20% of means, (ii) lowering stimulus duration by 20% resulted in a complete failure of uEPSCs, (iii) an increase in stimulus duration by 20% neither changed mean amplitude nor shape of uEPSCs. To elicit AMPAR- and NMDAR-mediated components of uEPSCs, records were performed at holding potentials of −70 mV and + 50 mV, respectively.

### Synaptic EAAT2 immunofluorescence

Using deep isoflurane anesthesia, mice were transcardially perfused with ice-cold phosphate-buffered saline (PBS) followed by a solution of 4% (w/v) paraformaldehyde in PBS. Sagittal sections (30 µm) were prepared as previously described (Rothe et al., 2015). Freely floating sections were double-stained with guinea pig anti-vGluT1 (1:1000) and rabbit anti EAAT2 (1:2000), followed by respective secondary antibodies at a concentration of 1:800. Grey scale 16 bit images (1091 × 1091 pixels, pixel size 0.073 µm, no binning) were acquired from the dorsal striatum using a Zeiss 100x oil immersion objective (NA1.3) and a Spot Insight camera system (Diagnostic Instruments Inc, Michigan, USA). All images were taken from the dorsal striatum. Areas of interest (AOIs, 400 × 400 pixels, 853 µm^2^) were cropped from the larger viewfields, selecting neuropil areas with a minimum of cell somata or vessels. Quantification of EAAT2 immunofluorescence (IF) was performed using ImagePro Plus (MediaCybernetics, Roper, Sarasota, USA). Within the selected AOIs, smaller regions of interest (ROIs, 25 × 25 pixels, 3.33 µm^2^) were then centred to individual vGluT1+ spots to determine the level of synaptic EAAT2 IF. A threshold algorithm was used to define the boundaries of the EAAT2+ area excluding pixels with F < ROI mean + 1.5 SD. The data is expressed as integral intensity of suprathreshold pixels. The term “*Synaptic integral EAAT2 IF*” refers to the mean value from 10 individually assessed ROIs (i.e. the environment of 10 vGluT1+ terminals) within one AOI. The sections from 3 WT and 3 Q175 HOM were stained together, and all images were acquired with the same camera settings. A total of 300 synapses were evaluated per genotype.

### Statistics

Data analysis was performed with Prism 8 (GraphPad, San Diego, USA). Considering that the comparison of the means could be influenced by inter-animal variance (Aarts et al., 2014) we have performed multi-level (“nested data”) analysis, where needed. P values of <0.05 were considered statistically significant. Significance levels were marked by asterisks, where * corresponds to P < 0.05, ** - P < 0.01 and *** - P < 0.001. The numbers indicate animals, cells or presynaptic terminals, as mentioned in the figure legends or tables. Genotype-related effects are described in % of WT levels (Δ of Tabs. 2, 3) or as effect strength according to Cohen’s D or Hedges’ G. D or G values larger 0.8 suggest that the respective effect was strong.

## RESULTS

### Evaluation of action potential-(AP-)mediated perisynaptic corticostriatal Glu transients using the new ultrafast sensor iGlu*_u_*in acute slices from adult mice

Placement of stimulating electrodes in the vicinity of corticostriatal terminals at rest was carried out under visual guidance (Fig. 1A). Bouton size was defined on the basis of resting fluorescence in the region of interest, ROI (Fig. 1, thick blue outline). The deduced virtual *Bouton diameter* exhibited a bimodal distribution (Fig. 1C). Varicosities with a diameter <=0.57 µm were defined as “Small” and, for the sake of brevity, tentatively referred to as IT type. Accordingly, varicosities with d >=0.63 µm were classified as “Large” or PT type. The size difference between terminals classified as Small (IT) vs. Large (PT) was significant at P < 0.001 (Tab. 1).

**Tab. 1.** Comparison of iGlu_u_ signals in varicosities type “Small” (presumably IT) and “Large” (presumably PT). Peak amp – Peak amplitude: ΔF/F at the peak of averaged transient derived from all suprathreshold pixels. TauD – time constant of decay derived from fitting a monoexponential function to the decay from peak amplitude. Peak spread – peak of the spread transient. See Methods section for more details. N-t – number of terminals. N-a – number of animals. Δ(%) – difference to WT in % of WT (=100%). In bold: indicators with significant afferent-related difference according to multi-level analysis (terminals nested in animals).

| iGlu<sub>u</sub> signals at IT vs. PT terminals | Varicosities type "Small" |  |  |  | Varicosities type "Large" |  |  |  | Statistics (nested t test) |  |  |
| --- | --- | --- | --- | --- | --- | --- | --- | --- | --- | --- | --- |
|  | Mean | SE | N-t | N-a | Mean | SE | N-t | N-a | F (DFn,Dfd) | P | Hedge's G |
| **Bouton diameter (μm)** | **0.51** | **0.01** | **12** | **6** | **0.74** | **0.02** | **20** | **3** | **77.00 (1,14)** | **<0.001** | **4.018** |
| Peak amp #1 (%) | 22.72 | 3.22 | 12 | 6 | 35.97 | 6.02 | 20 | 10 | 2.30 (1,14) | 0.152 | 0.590 |
| TauD #1 (ms) | 4.65 | 0.75 | 12 | 6 | 4.56 | 0.39 | 20 | 10 | 0.01 (1,30) | 0.906 | 0.044 |
| Peak spread #1 (μm) | 1.54 | 0.19 | 12 | 6 | 1.24 | 0.11 | 20 | 10 | 1.76 (1,14) | 0.206 | 0.518 |
| **First/last ratio peak amp, 20 Hz** | **0.68** | **0.06** | **12** | **6** | **1.18** | **0.05** | **20** | **10** | **37.50 (1,14)** | **<0.001** | **2.382** |
| **First/last ratio peak amp, 100 Hz** | **0.98** | **0.09** | **10** | **6** | **1.61** | **0.14** | **17** | **9** | **10.1 (1,25)** | **0.004** | **1.269** |
| **First/last ratio peak spread, 20 Hz** | **0.74** | **0.04** | **12** | **6** | **1.02** | **0.05** | **20** | **10** | **11.9 (1,14)** | **0.004** | **1.367** |
| **First/last ratio peak spread, 100 Hz** | **1.13** | **0.1** | **10** | **6** | **1.5** | **0.08** | **17** | **9** | **7.58 (1,25)** | **0.011** | **1.097** |
| **Integral ΔF/F #1-6, 100 Hz** | **0.89** | **0.14** | **10** | **6** | **2.28** | **0.51** | **17** | **9** | **6.05 (1,12)** | **0.032** | **0.828** |
| **Residual ΔF/F amp after #6/peak amp #1, 100 Hz** | **0.38** | **0.1** | **10** | **6** | **0.77** | **0.1** | **17** | **9** | **6.47 (1,25)** | **0.018** | **1.013** |
| **Residual ΔF/F spread after #6, 100 Hz (μm)** | **1.05** | **0.15** | **10** | **6** | **1.66** | **0.15** | **17** | **9** | **5.21 (1,12)** | **0.042** | **1.071** |

After electrical stimulation of a fluorescent corticostriatal axon in the dorsal striatum iGlu*_u_* intensity increased in the pixels adjacent to the bouton at rest (Fig. 1D, thin black outline: active area). To assess the dynamic characteristics of the iGlu*_u_* signal, the mean values of all supra-threshold pixel intensities (ΔF/F) generated by one synapse were plotted against time (Fig. 1E). The *Peak amplitude* was determined, and a monoexponential function (red line) was fitted to the averaged iGlu*_u_* transient ΔF/F to determine *TauD*. In the case of single pulse activation, there was no significant correlation between *TauD* and *Bouton diameter* (not illustrated).

The focus of the current experiments was placed on the time course of the iGlu*_u_* signals. The position of the sensor and its low affinity for Glu naturally set limits to the detection of [Glu] elevations at larger distance from the site of vesicle exocytosis. Nevertheless, we also expected some preliminary information on the spatial characteristics of the iGlu*_u_* signal. Therefore, the parameter *Peak spread* was deduced from the projection of the supra-threshold iGlu*_u_* area to the focal plane and plotted against time (Fig. 1F, distance between dotted red lines). Under condition of single pulse activation, *Peak spread* exhibited a significant positive correlation with *TauD* (Fig. 1G), but there were no terminal-type-related differences in the mean values of *Peak amplitude*, *TauD* and *Peak spread* after single pulse activation (Fig. 1H, Tab. 1).

The contrasting properties of IT- and PT-type terminals became more obvious with repeated stimulation. Activation with stimulus pairs at an interval of 50 ms revealed differences in the paired pulse ratio (PPR) of *Peak amplitude* resulting in a positive correlation between PPR and *Bouton diameter* (Fig. 1I). This finding validated our size criterion for synapse identification and provided a first hint that IT and PT afferents may generate a differential load for Glu clearance when repeatedly activated.

### Frequency-dependent potentiation of Glu release at PT but not IT corticostriatal terminals

As we aimed at exploring the limits of Glu release under conditions resembling the cortical activity during movement initiation, we applied 2 or 6 stimuli at frequencies of 20 or 100 Hz to elicit AP-mediated Glu release. At all frequencies tested, Small/IT and Large/PT terminals exhibited contrasting types of short-term plasticity, i.e. depression or no change in IT and potentiation in PT terminals (Fig. 2A-D and Tab. 1). The *Peak amplitude* observed after the last stimulus in a train normalized to the response #1 were larger in PT than in IT terminals (Fig. 2E, Tab. 1). The normalized *TauD* values exhibited little difference (Fig. 2F), but the normalized *Peak spread* differed, being larger at PT-type varicosities (Fig. 2G). When tested at 100 Hz, PT synapses produced larger *Integral ΔF/F #1-6* than IT terminals (Fig. 2H) and accumulated more *Residual ΔF/F* (Fig. 2I). The data suggests that the stimulus-locked response to the last AP adds to already incompletely cleared synaptic Glu. Thus, under repetitive activation conditions, corticostriatal afferents might be affected by conditions of weak astrocytic Glu uptake.

### Directly induced Glu transients in tetrodotoxin (TTX)

With the AP mechanism intact, fluorescence might also originate from neighbouring release sites, especially if the axon heads deeper into the z-plane of the slice. In case of the serial (*en passant*) type synapses (as characteristic of PT afferents) this could erroneously increase signal duration and spread. Another caveat to be faced in the case of HD preparations is a possible alteration of voltage-activated channels in the cortical afferents (Silva et al., 2017) which may affect the duration of the presynaptic depolarization, the influx of Ca^2+^ and, consequently, the amplitude and duration of iGlu*_u_* signals, without having a direct impact on the clearance machinery of the astrocytes. Moreover, respective deficits might preferentially occur in IT or PT axons. Considering these complexities it was decided to by-pass the AP mechanism by directly depolarizing the glutamatergic terminals in TTX and to focus, initially, on just one type of terminal. We selected the Large/PT input.

To achieve in TTX [Glu] elevations similar to those obtained under physiological activation conditions from PT terminals at 100 Hz, it was sufficient to increase [Ca^2+^]_ec_ to 5 mM and to add a hyperpolarizing prepulse to the standard 1 ms depolarization used both in AP and TTX experiments. In the absence of a conditioning hyperpolarizing pre-pulse, the direct depolarization was insufficient to elicit release (Fig. 3A). The results of Fig. 3B, C indicate that the selected protocol provided a good match between the physiologically induced #6 responses at 100 Hz and the directly induced responses in TTX. In any case, iGlu*_u_* elevations were completely abolished by the Ca^2+^ channel blocker Cd^2+^ (Fig. 3D). This stimulation protocol was then expected to provide a reasonably standardized challenge of the synaptic Glu uptake in WT or HD mice. In the following experiments (Fig. 4 and 5) all synapses were tested in TTX applying paired (Δt = 50 ms) biphasic pulses with a repetition frequency of 0.1 Hz.

**Fig. 3.**
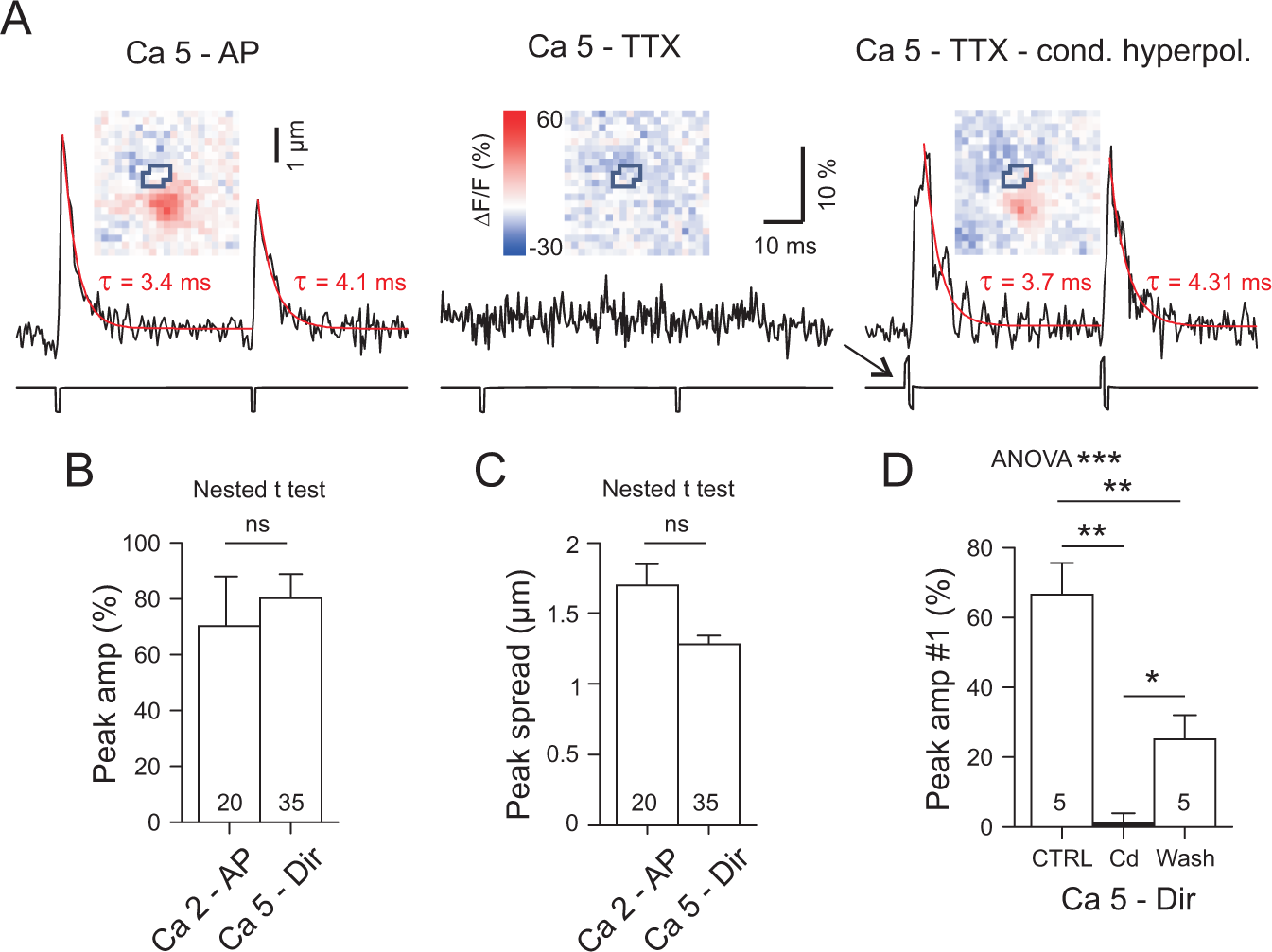
Responses to directly induced test pulses by-passing the AP mechanism in WT boutons of type PT. (A) Experiment in elevated [Ca^2+^]_ec_. Left: Response to electrical stimulation in the absence of TTX elicited by a short (1 ms) depolarizing pulse. In 5 mM Ca^2+^ the AP-mediated response exhibits paired pulse depression. Middle: same condition but in the presence of TTX. Note complete block of Glu release. Right: Response in 5 mM Ca^2+^ and TTX, but elicited with the 1 ms depolarizing pulse preceded by a short hyperpolarizing pulse. This configuration will in the following be referred to as “Ca 5 - Direct”. (B, C) Stimulus intensity for direct activation of Glu release in TTX (“Dir - Ca 5”) was adjusted such that the peak amplitude and peak spread of iGlu_u_ signals matched the amplitudes observed with the last (#6) 100 Hz response under physiological conditions (“AP - Ca 2”). (D) The directly induced responses were completely blocked by Cd^2+^ (500 µM). Cohen’s D: CTRL vs. Cd = 4.4, Cd vs. Wash = 2, CTRL vs. Wash = 2.3. * - P < 0.05, ** - P < 0.01, *** - P < 0.001.

**Fig. 4.**
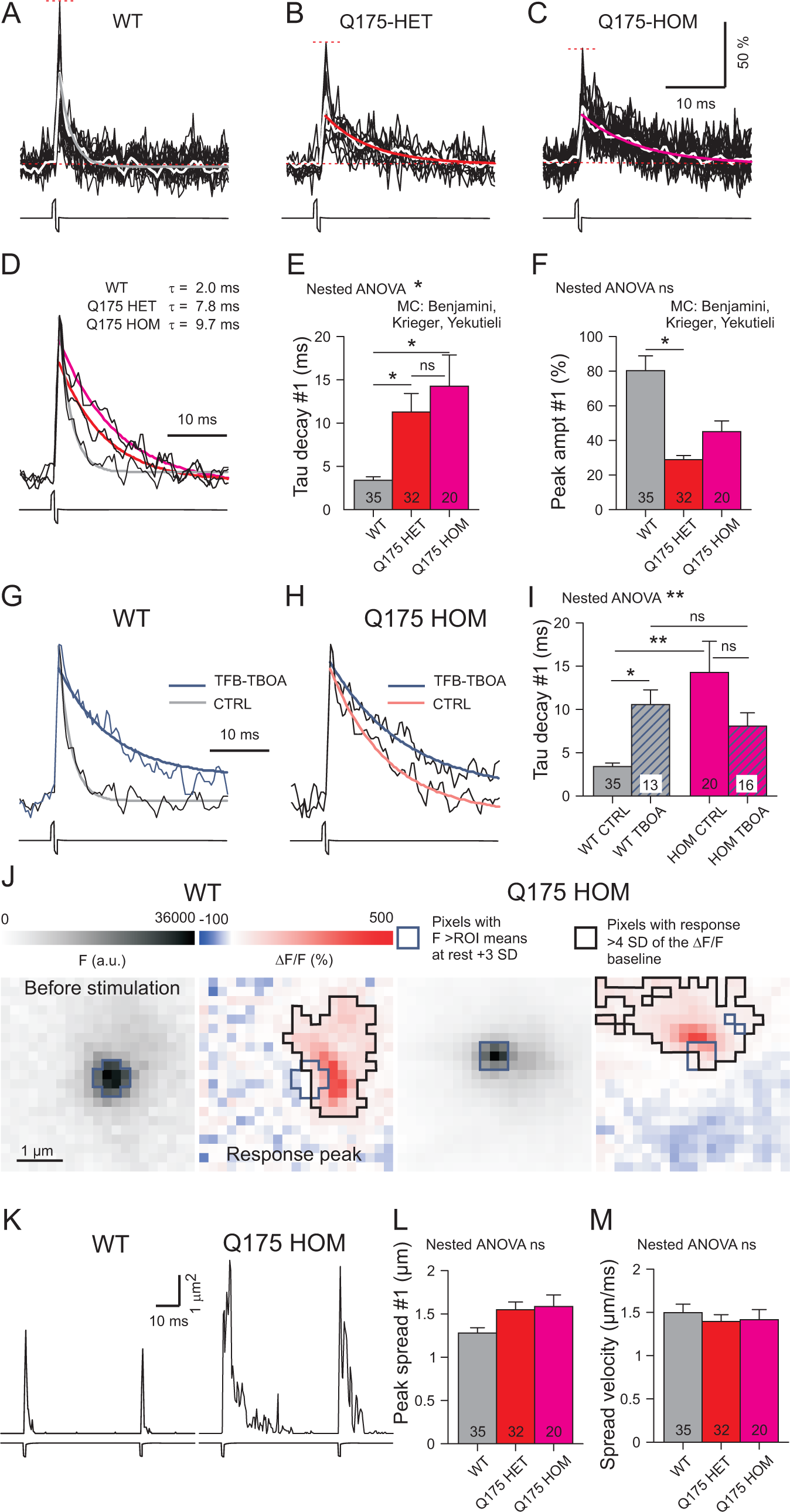
HD-related differences in the clearance of synaptically released Glu. (A-C) Superposition of suprathreshold pixel transients induced by direct activation of PT-type varicosities in the the presence of TTX. Differences between dotted red horizontal lines: Maximal amplitude of a single pixel transient. In white: averaged transient from all suprathreshold pixels. Curve in grey (WT), red (HET) and magenta (HOM) - monoexponential function fitted to the decay from peak amplitude. (D) Averaged responses normalized to same peak amplitude, same boutons as in in (A-C). The respective fitting curves highlight the differences in the duration of the Glu transients. (E, F) Quantification of the results from the entire data set. WT - gray, HET - red and HOM - magenta. (G) Incubation of WT slices in 100 nM of TFB-TBOA simulated the depression of Glu clearance observed in HOM (H, I) (J-L) Specimen images, traces and quantification for the spread in WT and HD mice. (M) Lack of genotype-related differences in spread velocity. Nested ANOVA statistics: (E) F(2,26) = 4.17, P = 0.027. (F) F(2,26) = 2.7, P = 0.086. (I) F(3,15) = 5.5, P = 0.0095. (L) F(2,26) = 0.52, P = 0.600. (M) F(2,26) = 0.65, P = 0.528. * -

**Fig. 5.**
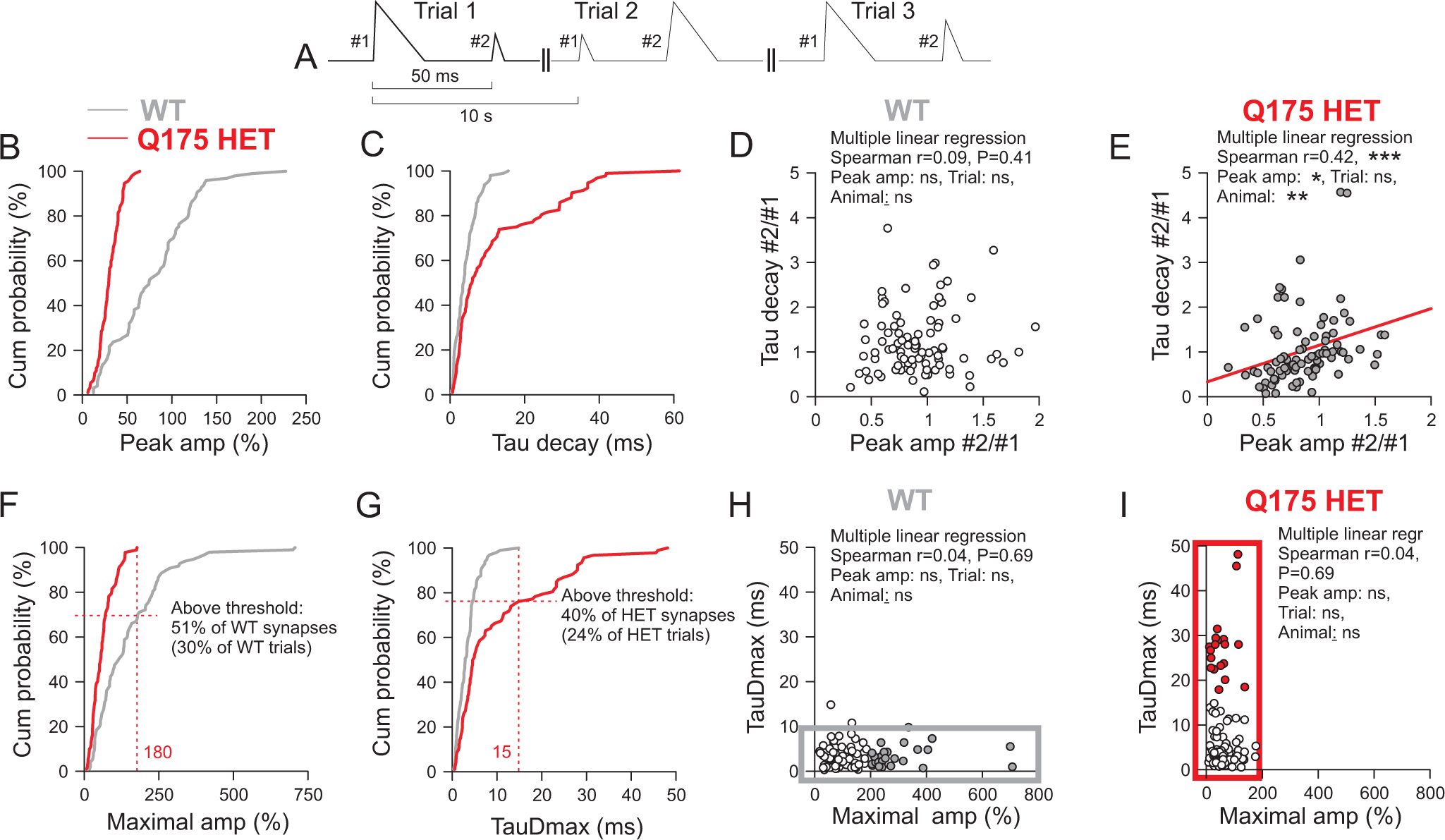
Relationship between Glu release and uptake and identification of dysfunctional synapses in HD. (A) Basal scheme of data organization. The graphs of (B-I) are based on 3 consecutive trials from each synapse. Data from 35 PT-type WT synapses and 32 HET synapses, except (D, E). (B, C) Cumulative histograms of #1 Peak amplitude and #1 TauD values. (D, E) Plots of normalized (to #1of the first trial) #2 responses. Data from 31 PT-type WT synapses and 30 HET synapses. Note that HET but not WT exhibited a positive correlation between TauD and Peak amplitude. In (E) the slope parameters of the three predictors Peak amplitude, Trial and Animal were significant for Peak amplitude (P = 0.015) and Animal (P = 0.002). The latter suggests that in different Q175 HET the disease has progressed to different degree. Significance levels for other variables: Peak amp*Animal -P = 0.003, Peak amp*Trial - n.s., Trial*Animal - n.s. (F, G) Cumulative histograms of #1 Maximal amplitude and #1 TauDmax. In the HET sample all Maximal amplitude values were < = 180%. In the WT sample all TauDmax values were <=15 ms. A total of 40 synapses exhibited in at least 1 of the 3 trials a TauDmax value exceeding the Threshold defined by the longest TauDmax in WT. 24% of the HET trials exceeded the 15 ms limit. Accordingly 30% of the WT responses were larger than in HET, and these suprathreshold responses were derived from 51% of the synapses. (H, I) Correlograms of TauDmax and Maximal amplitude for WT and Q175 HET. These graphs emphasize the HD-related differences in the ranges of Maximal amplitude and TauDmax. Maximal amplitude values exclusively seen in WT are shown in grey, and TauDmax values exclusively encountered in HET are shown in red. * - P < 0.05, ** - P < 0.01, *** - P < 0.001.

### Slowed Glu clearance at single PT-type corticostriatal terminals in HD

In the Q175 mouse model of HD, motor symptoms (hypo- and dyskinesia, pathological circling) develop quite slowly. However, at the age of one year and older, both Q175 HET and HOM resemble the human phenotype at a symptomatic stage (Khakh et al., 2017). In Q175 HOM motor impairment coincided with the appearance of pathological gamma oscillations in the local field potential (LFP) recordings at quiet rest (Rothe et al., 2015). In R6/2 mice, the changes in the LFP power spectrum were less pronounced after treatment with ceftriaxone (CEF), a transcriptional activator reported to increase the level of EAAT2 protein in the dorsal striatum (Miller et al., 2012). TFB-TBOA is a blocker of Glu uptake (Shimamoto et al., 2004). Its application would therefore simulate the effect of reduced EAAT2 expression/activity in HD.

Fig. 4A-C shows representative single synapse records from WT, HET and HOM. The black traces are individual pixel transients. The values between the dotted lines correspond to the *Maximal amplitude*. The white line is the mean transients derived from all suprathreshold pixels of a synapse. The monoexponential fitting curves are shown in grey (WT), red (HET) or magenta (HOM). Fig. 4D presents the amplitude-scaled average responses for the three genotypes illustrating our main finding: In HD slices the iGlu*_u_* transients decay more slowly than in WT. Interestingly, this HD-related alteration was significant not only in Q175 HOM but also in HET (Fig. 4E) thereby demonstrating the usefulness of the Q175 HET model for research on astrocyte pathology in HD.

An important additional question concerns the amount of released Glu. Is it increased by HD?- This was not the case, on the contrary. Despite a considerable variability in the *Peak amplitude*, we found a significant difference between WT and Q175 HET (Fig. 4F, Tab. 2). Unfortunately, multilevel data analysis failed to verify the difference between WT and HOM, due to the small number of available HOM. Our result is, however, in line with the data from R6/2 (Parievsky et al., 2017) suggesting that the presently disclosed HD-related prolongation of the iGlu*_u_* signal occurs despite a concomitant decrease in the Glu output from single PT terminals.

**Tab. 2.**
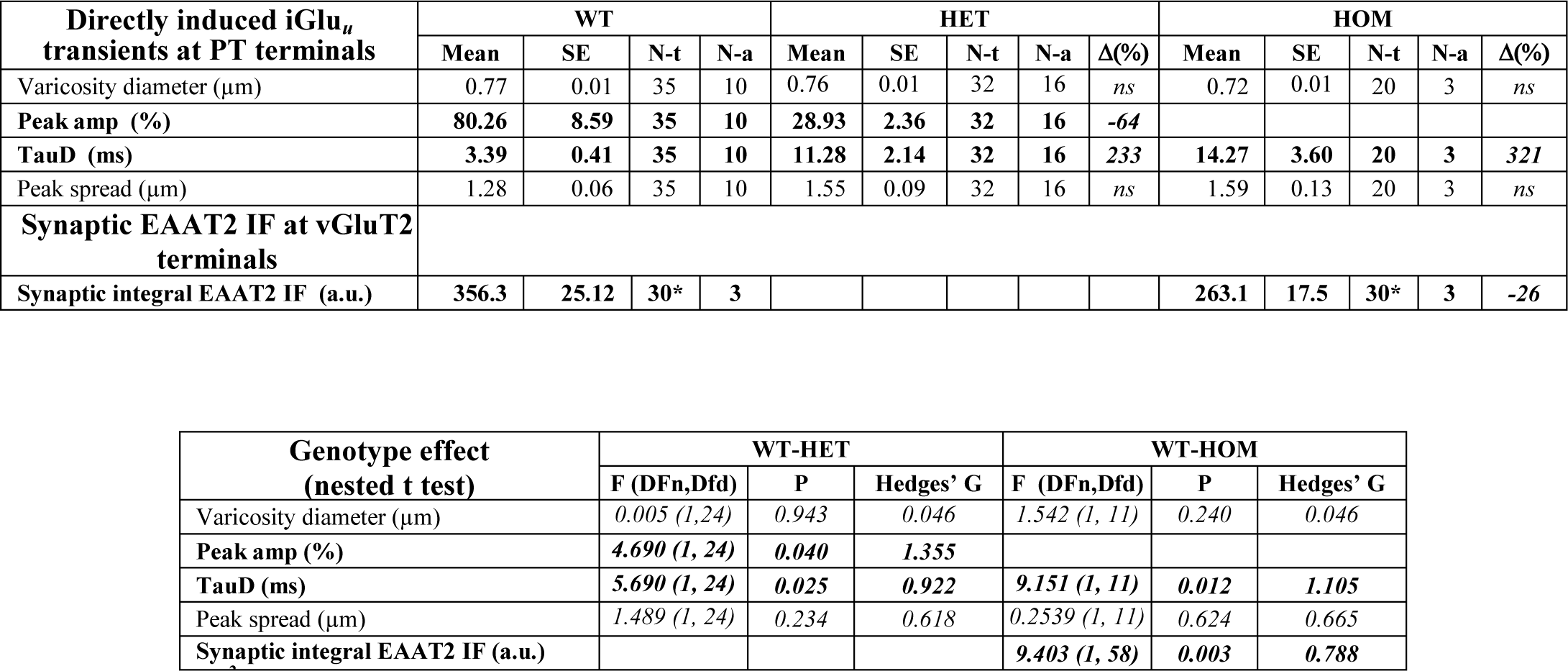
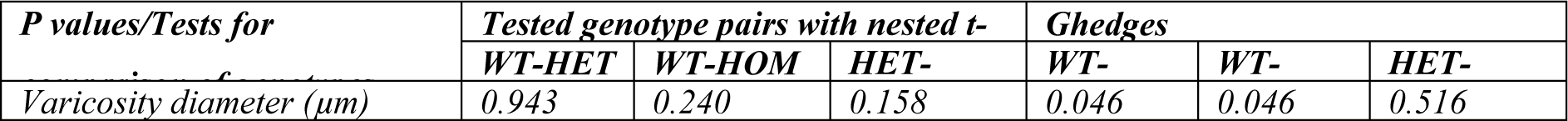
Comparison of WT with Q175 HET or HOM. Peak amp – peak amplitude: ΔF/F at the peak of averaged transient derived from all suprathreshold pixels. TauD – time constant of decay derived from fitting a monoexponential function to the decay from peak amplitude. Peak spread – peak of the spread transient. See Methods section for more details. N-t – number of terminals. N-a – number of animals. Δ(%) – difference to WT in % of WT (=100%). *Each data point represents the mean value from 10 synapses within one area of interest. In bold: indicators with significant afferent-related difference according to multi level analysis (level 1: animals, level 2: terminals).

| Directly induced iGlu<sub>u</sub> transients at PT terminals | WT |  |  |  | HET |  |  |  |  | HOM |  |  |  |  |
| --- | --- | --- | --- | --- | --- | --- | --- | --- | --- | --- | --- | --- | --- | --- |
|  | Mean | SE | N-t | N-a | Mean | SE | N-t | N-a | Δ(%) | Mean | SE | N-t | N-a | Δ(%) |
| Varicosity diameter (μm) | 0.77 | 0.01 | 35 | 10 | 0.76 | 0.01 | 32 | 16 | *ns* | 0.72 | 0.01 | 20 | 3 | *ns* |
| Peak amp (%) | **80.26** | **8.59** | **35** | **10** | **28.93** | **2.36** | **32** | **16** | **-64** |  |  |  |  |  |
| TauD (ms) | **3.39** | **0.41** | **35** | **10** | **11.28** | **2.14** | **32** | **16** | **233** | **14.27** | **3.60** | **20** | **3** | **321** |
| Peak spread (μm) | 1.28 | 0.06 | 35 | 10 | 1.55 | 0.09 | 32 | 16 | *ns* | 1.59 | 0.13 | 20 | 3 | *ns* |
| Synaptic EAAT2 IF at vGluT2 terminals |  |  |  |  |  |  |  |  |  |  |  |  |  |  |
| Synaptic integral EAAT2 IF (a.u.) | **356.3** | **25.12** | **30\*** | **3** |  |  |  |  |  | **263.1** | **17.5** | **30\*** | **3** | **-26** |

**Tab. 2. Comparison of WT with Q175 HET or HOM.**
| Genotype effect<br>(nested t test) | WT-HET |  |  | WT-HOM |  |  |
| --- | --- | --- | --- | --- | --- | --- |
|  | F (DFn, Dfd) | P | Hedges' G | F (DFn, Dfd) | P | Hedges' G |
| Varicosity diameter (μm) | 0.005 (1, 24) | 0.943 | 0.046 | 1.542 (1, 11) | 0.240 | 0.046 |
| Peak amp (%) | **4.690 (1, 24)** | **0.040** | **1.355** |  |  |  |
| TauD (ms) | **5.690 (1, 24)** | **0.025** | **0.922** | **9.151 (1, 11)** | **0.012** | **1.105** |
| Peak spread (μm) | 1.489 (1, 24) | 0.234 | 0.618 | 0.2539 (1, 11) | 0.624 | 0.665 |
| Synaptic integral EAAT2 IF (a.u.) |  |  |  | **9.403 (1, 58)** | **0.003** | **0.788** |

| *P* values/Tests for | Tested genotype pairs with nested t- |  |  | Ghedges |  |  |
| --- | --- | --- | --- | --- | --- | --- |
|  | WT-HET | WT-HOM | HET- | WT- | WT- | HET- |
| Varicosity diameter (μm) | 0.943 | 0.240 | 0.158 | 0.046 | 0.046 | 0.516 |

The above observations do not immediately prove that the prolongation of the iGlu*_u_* transients in HD were due to altered functionality of the astrocytes. It was at least necessary to clarify whether *TauD* responded to pharmacological manipulation of astrocytic Glu transport. This was the case. The iGlu*_u_* transients of PT terminals exhibited a clear sensitivity to TFB-TBOA (Fig. 4G). In WT, 100 nM of the antagonist prolonged the iGlu*_u_*decay to the same extent as the disease (Fig. 4I), the effect of pharmacological EAAT2 block being less pronounced in Q175 HOM (Fig. 4H, I).

According to the presently available models of glutamatergic synapses (Zheng et al., 2008; Scimemi and Beato, 2009), a spread of >1.25 µm should be sufficient to activate extrasynaptic NMDA receptors. Although iGlu*_u_* expression in the presynaptic terminals cannot provide exhaustive information on the spatial characteristics of perisynaptic [Glu], we nevertheless examined the *Peak spread* (black outlines in Fig. 4J). Although there was a tendency for increase (Fig. 4K) this tendency failed to reach significance in Nested ANOVA (Fig. 4L) and nested t tests (Tab. 2). The mean spread velocity (about 1.5 µm/s) did not vary with the genotype (Fig. 4M).

### Positive correlation between Glu release and clearance in HD but not WT synapses

Cumulative histograms and correlograms were plotted for further analysis of HD-related synapse pathology. The graphs of Fig. 5, except (D, E), are based on the values obtained from 35 WT and 32 HET synapses, as explained by the evaluation scheme in *(A)*. Each synapse is represented with 3 consecutive trials elicited at a frequency of 1/10 Hz. The interval between the stimuli for #1 and #2 was 50 ms. All data is from experiments in TTX. *Fig. 5B*, C shows the relative probability of occurrence of *Peak amplitude* and *TauD*, based on a total of 105 #1 responses from 35 WT synapses and 96 #1 responses from 32 HD synapses.

Due to the highly variable configuration of the individual synapses with respect to the surrounding tissue and the focal plane of the camera, and due to inter-animal variation, the values obtained from the averaged iGlu*_u_* trials of different synapses exhibited considerable variability. It was therefore necessary to normalize the data. Among several possibilities, we chose the #1 response of every trial to normalize the #2 responses. Typically #2 responses were smaller after larger #1 responses, and *vice versa* (Fig. 5A). The normalization reduced the impact of inter-synapse variability in favour of inter-trial variability. If in a given trial glutamate output touched the limits of uptake one could expect that such release event would produce a prolonged [Glu] transient. In contrast, if uptake capacity were sufficient for any amount of released glutamate, the fluctuating *TauD* values should be independent on *Peak amplitude*. It can be seen (Fig. 5D, E) that HD but not WT synapses displayed a positive correlation between *TauD* and *Peak amplitude*, consistent with the proposal that in HD some synapse exhibited signs of clearance insufficiency, the prolongation of *TauD* being more pronounced in trials with enhanced *Peak amplitude*.

### Identification of dysfunctional synapses

The pixels with the highest stimulus-induced elevations of ΔF/F were always located within or immediately next to the boundaries of the resting terminal (see Fig. 1D). The *Maximal amplitude* derived from the highest single pixel transient can be regarded as a measure of the Glu output while *TauDmax* would reflect the clearance at the site of release minimizing the influence of Glu diffusion. *Fig. 5F*, G presents the cumulative probabilities of occurrence of *Maximal amplitude* and *TauDmax*. One can see that none of the HET entries of *Maximal amplitude* were larger 180%, and about 24% of *TauDmax* entries exceeded 15 ms. 40% of the tested synapses generated TauDmax in at least 1 of the 3 trials, and all of these responses were smaller than 180%. The differences in the ranges of these two indicators of release and clearance, respectively, are even more obvious in the correlograms of Fig. 5H, I. *TauDmax* values larger than 10 ms were (with 1 exception) absent in WT synapses tested with direct depolarization. As a first approximation, one can therefore state that, according to the distribution of *TauDmax* in WT and Q175 HET aged 15 to 19 months, 40% of HET synapses in the dosal striatum exhibited a pathological phenotype. Of course, this estimation is no more than an educated guess based on the assumptions that 15 ms is the largest TauDmax value to be expected in WT, and that in WT all synapses are fully functional. The data also hints that in dysfunctional synapses Glu may find its astrocytic transporter at bigger distance, in line with recent FRET data from corticostriatal synapses in R6/2 (Octeau et al., 2018).

### Reduced perisynaptic EAAT2 protein at corticostriatal terminals

To clarify whether the observed clearance deficit is indeed accompanied by a reduction of EAAT2 protein levels in the environment of corticostriatal terminals, we performed a quantification of EAAT2 IF in fixed sections, as described in the Methods and illustrated in Fig. *6A-M*.

**Fig. 6.**
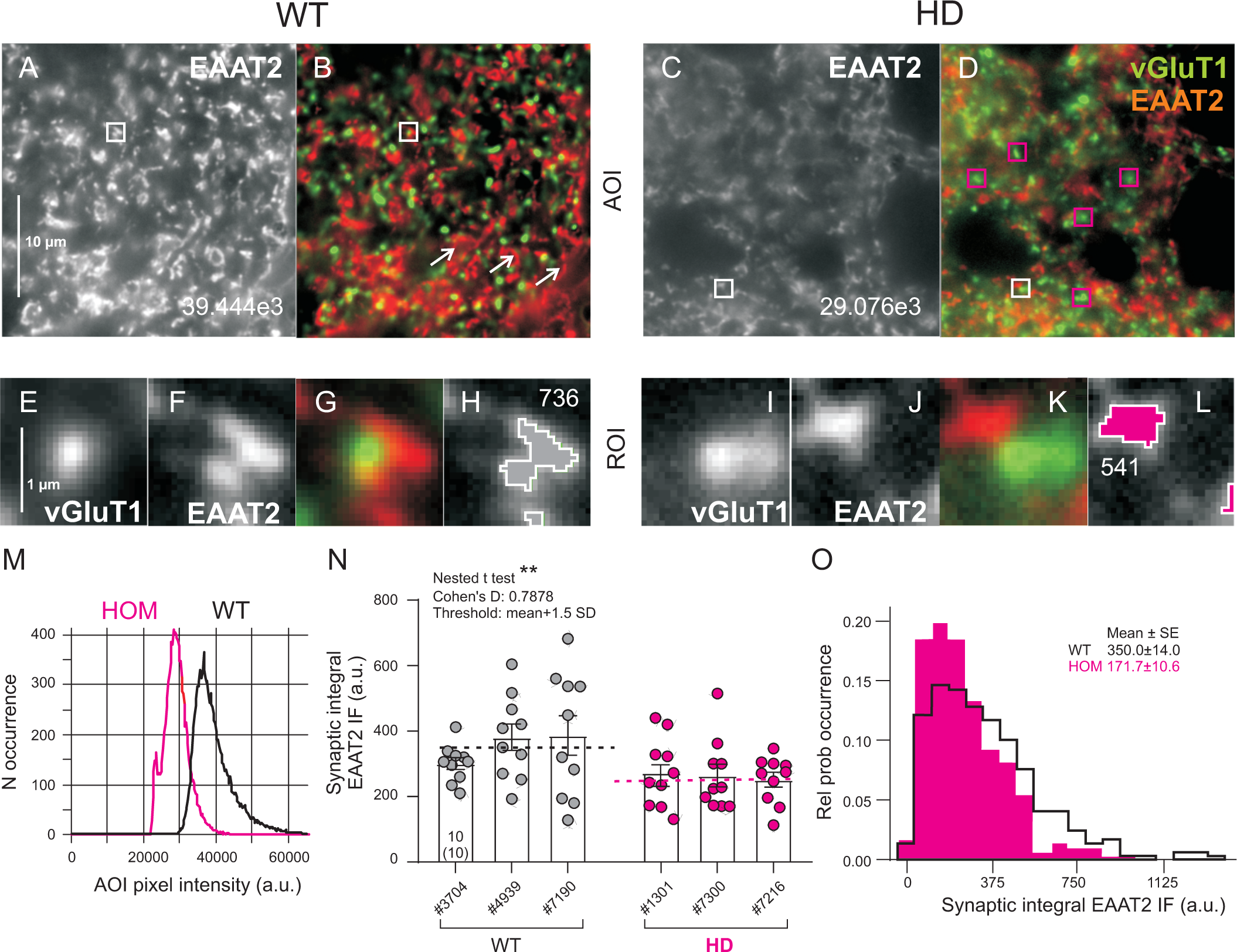
HD-related reduction of synaptic EAAT2 immunofluorescence (IF). Data from 3 male Q175 HOM (CAG range 176-191, age range 49-54 weeks) and 3 male WT siblings. (A, C) Areas of interest (AOIs) cropped from larger view fields for quantification of synaptic integral EAAT2 IF. Numbers on EAAT2 image: mean AOI IF intensity (no intensity threshold, same display range for WT and HOM). (B, D) Overlay of vGluT1 and EAAT2 images (display ranges optimized for object recognition). Squares outlined in white show ROIs as used for estimation of synaptic integral EAAT2 IF. Both WT and HOM images contain numerous EAAT2 clusters without synaptic terminals, presumably representing astrocytic end-feet in contact with blood vessels (arrows in B). In HD vGluT1 varicosities may occur without EAAT2 clusters (ROIs boxed in red). (E-H) and (I-L) enlarged ROIs showing (in this order) vGluT1, EAAT2, overlay and the suprathreshold EAAT2 as used for estimation of integral EAAT2 IF in the immediate vicinity of one corticostriatal vGluT+ terminals. Numbers on ROIs: integral suprathreshold fluorescence intensity for the EAAT2 channel. (M) Histogram of AOI EAAT2 pixel intensity. Note that “holes” from somata and blood vessels would influence the mean AOI values of EAAT2 intensity. N) Small ROI quantification of synaptic integral EAAT2 intensity by nested data analysis. Each data point represents the mean from 10 rectangular ROIs within one AOI. Dotted lines indicate mean level from 3 animals (with a total of 30 AOIs, 300 synapses) per genotype. Statistics (F, DFn, DFd): 9.403, 1, 58. P = 0.0033. Numbers on column: AOIs and animals (in brackets), same for all columns. (O) Histogram of synaptic integral EAAT2 from WT and Q175 HOM. n = 300 per group. Effect size (Cohen’s D) was obtained with the t-value calculated by the nested t test. Symbols, abbreviations: # animal number, a.u. arbitrary units. Color code: WT - light grey, HOM - magenta. * - P < 0.05, ** - P < 0.01, *** - P < 0.001.

It has frequently been observed, and could possibly be noticed in the examples of Fig. 6A-D, that in 100x images from the dorsal striatum of HD mice the areas preferentially occupied by neuropil (i.e. areas without “holes” from the somata of neurons and astrocytes) are smaller than in WT. Moreover, a variable fraction of vGluT1+ varicosities seemed to be devoid of synaptic EAAT2+ clusters, notably in HD (red boxes in Fig. *6D*). Finally, due to the presence of capillaries and the attached astrocyte end-feet, there were EAAT2+ clusters without vGluT+ counterparts (arrows in Fig. 6B). To avoid ambiguity resulting from these complexities, it was decided to quantify synaptic EAAT2 IF individually in sufficiently small ROIs (1.825 × 1.825 µm^2^) centred to just one vGluT1+-positive terminal. Fig. 6E-L shows representative ROIs selected from larger AOIs in the dorsal striatum (Fig. *6B*, D white boxes). A threshold algorithm was used to delineate the boundaries of the EAAT2 clusters from where the *Synaptic integral EAAT2 IF* values were actually sampled. Each data point in Fig. 6N represents the mean value from 10 ROIs of 1AOI. vGluT1+ terminals without any suprathreshold EAAT2 were avoided which may have caused an underestimation of the actual difference. Nested data analysis showed that the synaptic integral EAAT2 IF was significantly lower in HD (Fig. 6N, Tab. 2). Cohen’s D (0.7878) suggests a strong HD-related effect (- 26%). The histogram of synaptic integral EAAT2 intensity (Fig. 6O) illustrates the over-all shift towards lower values of *Synaptic integral EAAT2 IF* in individual ROIs.

### Prolonged NMDAR components of unitary EPSCs in HD

NMDARs are sensitive indicators of [Glu] and therefore well suited to detect a potentially existing Glu clearance deficits in the environment of active synapses, provided that the analysed responses are derived from one or few synapses only (Chiu and Jahr, 2017). CEF is known to stimulate the transcription of *SLC1A2*, i.e. the gene encoding EAAT2. It is therefore used to verify a contribution of EAAT2 in a pathology or recovery effect. Functional benefits from CEF injections have already been reported (Miller et al., 2008; Miller et al., 2012) and were attributed to enhanced EAAT2 expression in astroglia. Here we used focal optical stimulation of individual channel rhodopsin-expressing corticostriatal axons to record uEPSCs at −70 and + 50 mV. The experiments showed that the T50 value of the uEPSC recorded at + 50 mV is i) solely dependent on NMDARs, ii) prolonged in HD and iii) recovered to WT levels after treatment with CEF suggesting a sensitivity of corticostriatal input to the level of EAAT2 expression (Fig. 7A-C, Tab. 3). Other parameters of corticostriatal uEPSCs were found unchanged by HD (Fig. 7D-F). However, more work is needed to actually prove that the observed potentiation of NMDAR activity in striatal projection neurons had been a result of wider spread of synaptically released glutamate.

**Tab. 3.** Comparison of uEPSCs in WT, Q175 HOM and Q175 HOM treated with ceftriaxone (CEF). *Amplitude without failures. N – number of cells (c) or animals (a). MC – multiple comparison test according to Benjamini, Krieger, Yekutieli. Δ(%) - % change in comparison with WT (=100%). Note that the effect of genotype on TauD is both significant and strong (bold row).

| Corticostriatal unitary EPSCs | WT |  |  |  | HOM |  |  |  |  | HOM+CEF |  |  |  |  | Nested MC |  | Nested ANOVA |
| --- | --- | --- | --- | --- | --- | --- | --- | --- | --- | --- | --- | --- | --- | --- | --- | --- | --- |
| | Mean | SE | N | | Mean | SE | N | | $\Delta$% | Mean | SE | N | | $\Delta$% | WT/HOM | HOM/CEF | F/P/Hedges' G |
|  |  |  | c | a |  |  | c | a |  |  |  | c | a |  |  |  |  |
| Amp* NMDAR resp at +50 mV (pA) | 16.03 | 1.79 | 24 | 6 | 21.97 | 1.83 | 20 | 4 | - | 13.85 | 2.32 | 11 | 3 | - | 0.3202 | 0.1742 | 1.15/0.3533/0.700 |
| **T50 of NMDAR resp at +50 mV (ms)** | **20.68** | **1.95** | **24** | **6** | **35.74** | **3.71** | **20** | **4** | **73** | **21.25** | **2.63** | **11** | **3** | - | **0.0124** | **0.0117** | **4.67/0.0340/1.141** |
| Amp* of AMPAR resp at -70 mV (pA) | 29.07 | 2.82 | 27 | 6 | 38.71 | 2.54 | 27 | 5 | - | 30.05 | 3.03 | 26 | 5 | - | 0.1314 | 0.1016 | 1.65/0.2100/0.691 |
| TauD AMPAR resp at -70 mV (ms) | 4.06 | 0.24 | 27 | 6 | 4.76 | 0.31 | 27 | 5 | - | 4.05 | 0.29 | 26 | 5 | - | 0.1417 | 0.1111 | 1.84/0.1767/0.486 |
| CV of AMPAR resp at -70 mV (pA) | 42.55 | 3.78 | 27 | 6 | 31.21 | 2.80 | 27 | 5 | - | 37.78 | 2.79 | 26 | 5 | - | 0.1106 | 0.5351 | 1.35/0.2742/0.656 |
| Failure rate AMPAR resp at -70 mV (ms) | 8.14 | 1.51 | 27 | 6 | 4.91 | 1.01 | 27 | 5 | - | 6.37 | 1.29 | 26 | 5 | - | 0.1073 | 0.4196 | 1.51/0.2408/0.484 |

**Fig. 7.**
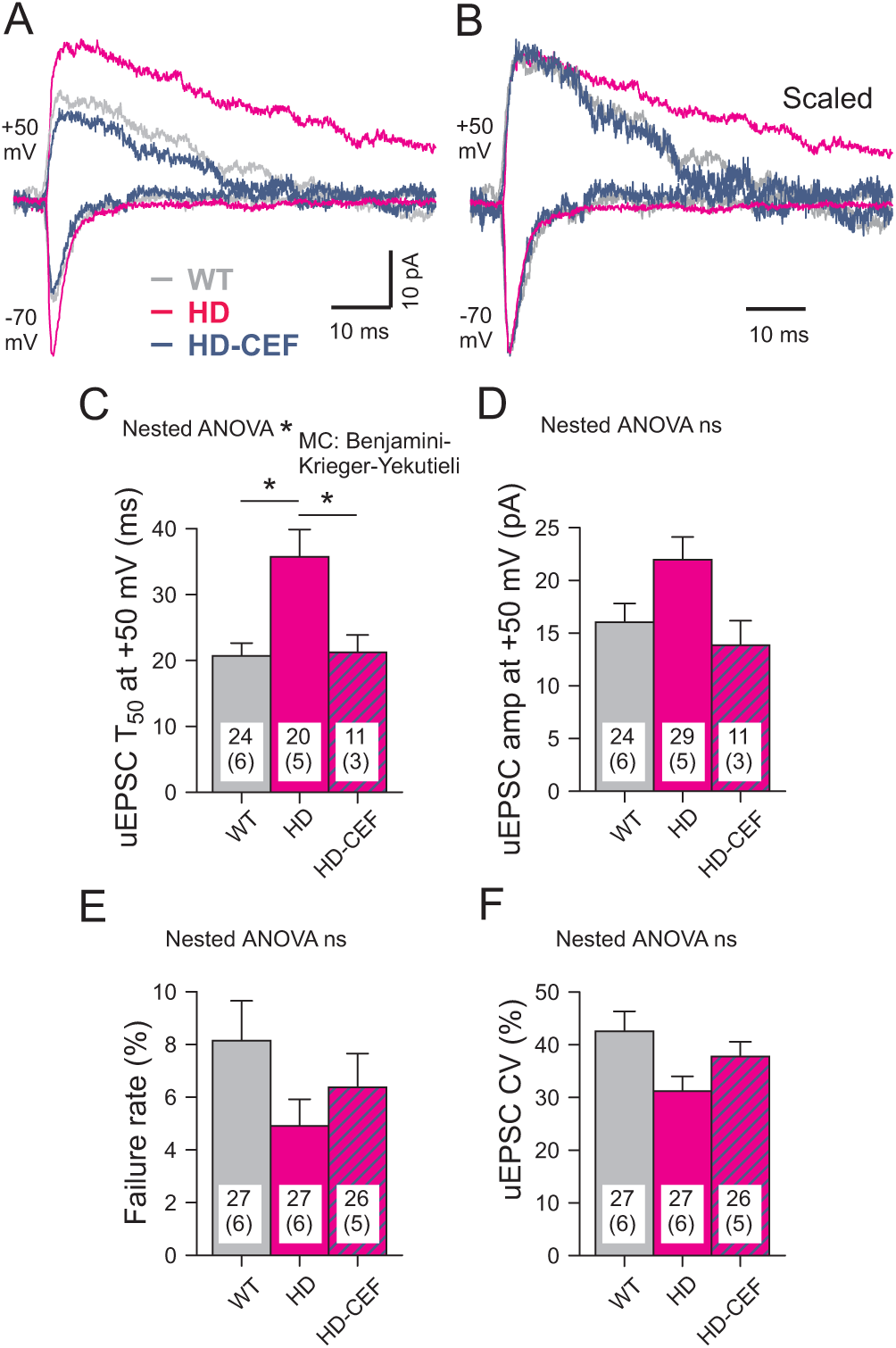
HD-related prolongation of the NMDAR component in uEPSCs elicited by optical stimulation of single corticostriatal afferents visualized by EYFP fluorescence after the expression of CaMKIIa.hChR2(E123 T/T159C)- EYFP.hGH. (A) Specimen traces as recorded at two different holding potentials in the presence of bicuculline methiodide (25 µM). (B) Same traces as (A) but aligned to peak. See prolonged decay in contrast to records at −70 mV. The half-decay time of the uEPSC (T50) was sensitive to APV (not illustrated). Treatment with CEF shortened the uEPSC at + 50 mV to WT level (blue traces). (C-F) Quantification of results. Note significantly larger T50 values of the NMDAR-mediated response at + 50 mV and recovery after CEF treatment (C). * - P < 0.05. Numbers in columns: number of tested SPNs and animals (in brackets). For detailed results of nested data analysis and Hedges’ G see Tab. 3.

## DISCUSSION

The analysis of single synapse iGlu*_u_*transients in acute slice preparations from adult mice provides new information on Glu clearance in its relation to the respective transmitter load. To summarize: 1) After single pulse activation, IT and PT synapses coped with the induced [Glu] elevations, but when challenged with high activation frequencies the [Glu] elevations produced by PT and IT terminals differed significantly (factor 2.6:1 for *Integral ΔF/F* at 100 Hz). 2) In HD, PT iGlu*_u_* transients were found to decay more slowly. About 40% of HD synapses (14/32 in HET can be regarded as deficient, considering the time needed for complete Glu clearance in any trial (TauD of iGlu*_u_* > 15 ms). 3) At any given terminal, the responses exhibited some inter-trial variability. Analysis of normalized #2 to #1 from 3 trials at the same synapse revealed that in Q175 HET, but not WT, iGlu*_u_* transients with larger *Peak amplitude* were associated with *larger TauD*. This is evidence for a disease-related loss of independence between the indicators of uptake and release. 4) HD decreased the range of *Maximal amplitude* but increased the range of *TauDmax*. 5) Immunostaining suggests that the immediate environment of corticostriatal terminals contains less EAAT2 protein. 6) The NMDAR-mediated unitary EPSCs elicited by optical stimulation of single ChR-expressing corticostriatal axons were prolonged in Q175 HOM.

### PT versus IT terminals

The unexpected differences in the properties of PT vs. IT terminals raise further questions on the mechanism(s) of release plasticity at the glutamatergic afferents to the dorsal striatum. Previous electrophysiological studies (Ding et al., 2008) implicated that corticostriatal connections preferentially exhibit paired pulse facilitation (PPF) while thalamostriatal connections are prone to paired pulse depression (PPD). Our results confirm preferential PPF with regard to the PT subgroup of corticostriatal afferents under physiological activation conditions (Tab. 1). However, as in any other synapse (for instance (Kirischuk et al., 2002)), a conversion from PPF to PPD is easily achieved by increasing the Ca^2+^ influx. There is a widely accepted rule of thumb suggesting that smaller initial responses are likely to produce facilitation, and *vice versa*. Considering that under the same experimental conditions PT and IT terminals produced about the same initial Glu output but opposite types of frequency-dependent plasticity, one can assume that these terminals indeed represent two classes of afferents with some differences in the presynaptic control of transmitter release.

Under condition of repetitive activation, the size of synaptic terminals and associated differences in the vesicle pool size could affect the integral Glu output, and also the degree of Glu escape (Genoud et al., 2006; Bernardinelli et al., 2014; Medvedev et al., 2014; Gavrilov et al., 2018). It has been hypothesized that thicker terminals could push the PAPs further away from the site of exocytosis which may result in wider signal spread if the transporters are challenged with a pronounced build-up of [Glu], as found in PT terminals.

### The hypothesis of non-saturating Glu uptake in healthy glutamatergic synapses

In view of a long history of changing opinions on the significance of astrocytic Glu transport as a possible determinant of synaptic strength it is good to have new tools at hand to shed light on the possible limits of Glu clearance in health and disease. Our uEPSC data from synaptic connections with one or few terminals confirm the long-standing idea that a weakness of Glu uptake has little influence on the decay kinetics of the fast desensitizing AMPA responses (Hestrin et al., 1990; Asztely et al., 1997; Goubard et al., 2011; Campbell et al., 2014). Moreover, our iGlu*_u_* data from healthy mice are in line with the more controversial prediction that in “normal” glutamatergic synapses glutamate transport would cope with any amount of physiologically released Glu (Diamond and Jahr, 2000; Tzingounis and Wadiche, 2007). Nevertheless, the present iGlu*_u_*-based postulate of non-saturating Glu uptake for IT- and PT-type corticostriatal synapses will need further verification under a wider range of conditions. It was already shown that the state of astrocytes could affect the structural plasticity of PAPs (Theodosis et al., 2008; Reichenbach et al., 2010; Bernardinelli et al., 2014; Heller and Rusakov, 2015; Verkhratsky and Nedergaard, 2018). Activity- and disease-dependent PAP retraction could produce a large variety of spill-out and spill-in effects which may not only change the access of the available transmitter(s) to respective neuronal and glial receptors, but also influence the efficacy of the astrocytic transport machinery itself (Armbruster et al., 2016).

### Evidence for impairment of Glu clearance in HD

Symptomatic HD is characterized by the loss of glutamatergic terminals in the dorsal striatum but it is still not clear whether this disease-related process of synapse pruning is to be attributed to glutamate excitotoxicity (Reiner and Deng, 2018). While the long-term consequences of reduced Glu clearance remain to be clarified, our present experiments provide new evidence suggesting that in symptomatic Q175 mice a significant fraction of PT (∼40%) synapses is afflicted by the disease, most likely exhibiting alterations in both uptake and release. When analysing the normalized *Peak amplitude* in 3 consecutive trials of the same synapse, it turned out that HD but not WT synapses displayed a positive, presumably pathological correlation between *Peak amplitude* and *TauD*. Considering in addition that i) treatment of WT with TBOA produced *TauD* values similar to those in HD, and ii) *Synaptic integral EAAT2 IF* was significantly less in HD, it is suggested that glutamate uptake, in general, and astrocytic EAAT2 deficiency, in particular, contribute to the observed synaptic dysfunction in HD.

However, this conclusion is not shared by all researchers. First of all, there is some evidence that EAAT2 is also localized on presynaptic terminals (Petr et al., 2013). In the R6/2 model of HD, the Rosenberg group confirmed the reduced expression of EAAT2 and the beneficial effects of CEF. But experiments with partial knock-down of *SLC1A2* revealed little change in the fraction of EAAT+ terminals and, even more important, in the progression of HD. Based on these and other findings, Rosenberg and colleagues questioned a role of EAAT2 in the pathogenesis of HD and forwarded the intriguing hypothesis that the observed down-regulation of EAAT2 applies to a nonfunctional intracellular fraction of the EAAT2 protein. We find the reported 40% reduction of the glutamate uptake activity in synaptosomes after conditional GLT1 knock-out (Petr et al., 2015) somewhat surprising considering that in the present material no more than 5% of the terminals exhibited full co-localization of vGluT1 and EAAT2 IF.

Considering the novelty of our present approach, it is not so unexpected that some results from other labs were not confirmed, in particular those obtained with the slow Glu indicator iGluSNFR (Marvin et al., 2013). (Parsons et al., 2016) activated glutamate release by high-frequency electrical field stimulation and used the iGluSnFR sensor to record Glu elevations in large viewfields. They found no HD-related difference in the fluorescence decay, in contrast to (Jiang et al., 2016). Both studies were carried out in R6/2 mice, the main difference being the site of expression of the Glu sensor (neurons vs. astrocytes). (Parievsky et al., 2017) applied optical field stimulation of channel-rhodopsin-(ChR2(H134R))- expressing corticostriatal axons to induce EPSCs in SPNs. This approach showed no increase in the decay times (T90-10) of NMDAR-mediated currents. On the contrary, the latter were significantly shorter in HD. However, considering the mean T90-10 values of this study (∼750 ms) it seems possible that the asynchrony of release characteristic produced by this type of optical field stimulation may not give the resolution needed for the estimation of synaptic Glu clearance. In general, space-and volume-averaging effects resulting from bulk activation of synaptic and non-synaptic Glu release and low resolution of the electrical or fluorescent signals can be expected to influence the interpretation of results on Glu uptake and release (see (Jensen et al., 2017; Reynolds et al., 2018) for a concise summary on these issues).

A question receiving growing attention in the field of synaptic plasticity and dysfunction is the role of other glutamate uptake mechanisms. Scimemi and colleagues (Bellini et al., 2018) illuminated the role of Glu uptake from two sides - pathology and functional rescue. Their convincing evidence suggests that the neuronal Glu transporter EAAT3 (EAAC1) ensures long-term synaptic activity by reducing the activation of mGluR1 in the striatum.

The ultimate proof of Glu uptake deficiency as a cause of synapse pathology in the dorsal striatum will be the recovery of normal synaptic performance after a therapeutic intervention targeting the astrocytes. Most intriguing, intrastriatal injection of a recombinant viral Kir4.1 vector restored a normal level of EAAT2 protein (Tong et al., 2014). However, it is not yet clear whether a mere stimulation of EAAT2 expression would suffice to achieve the desired reversal of motor symptoms in HD, because synaptic targeting and the activity of Glu transporters are also influenced by local translation (Sakers et al., 2017), lateral mobility (Murphy-Royal et al., 2015) and internalization (Leinenweber et al., 2011; Ibanez et al., 2016). Clearly, much more information is needed to fully understand the regulation of Glu uptake in the context of other astrocytic signaling cascades.

## ACKNOWLEDGEMENTS

We thank V. Beaumont, H. Kettenmann and S. Hirschberg for helpful discussions. D. Betances, A. Schönherr and J. Rösner provided skilled technical assistance. The work of the Grantyn lab was supported by CHDI (A-12467), the German Research Foundation (Exc 257/1) and intramural Charité Research Funds. Development of iGlu*_u_* in K. Török’s laboratory was funded by BBSRC grants BB/M02556X/1 and BB/S003894/1. N. Helassa is supported by a British Heart Foundation Intermediate Basic Science Research fellowship (FS/17/56/32925).

